# Evidence of Chelonid Herpesvirus 5 (ChHV5) in Green Turtles (*Chelonia mydas*) from Sabah, Borneo

**DOI:** 10.1101/2021.01.25.428031

**Authors:** Aswini Leela Loganathan, Pushpa Palaniappan, Vijay Kumar Subbiah

**Affiliations:** Biotechnology Research Institute, Universiti Malaysia Sabah, Kota Kinabalu, Sabah, Malaysia; Monash University Malaysia Genomics Facility, Monash University Malaysia, Selangor, Malaysia; Borneo Marine Research Institute, Universiti Malaysia Sabah, Kota Kinabalu, Sabah, Malaysia

**Author notes:** **Corresponding Author:** Vijay Kumar Subbiah, ^1^Biotechnology Research Institute, Universiti Malaysia Sabah, Jalan UMS, 88400, Kota Kinabalu, SABAH, MALAYSIA., Email address.

**Keywords:** Borneo, ChHV5, *Chelonia mydas*, Fibropapillomatosis, Mabul Island, Sabah, Malaysia

## Abstract

Fibropapillomatosis (FP) is characterized by cutaneous tumours and is associated with Chelonid herpesvirus 5 (ChHV5), an alphaherpesvirus from the family Herpesviridae. Here, we provide the first evidence of ChHV5-associated FP in endangered Green turtles (*Chelonia mydas*) from Sabah, which is located at the northern region of Malaysian Borneo. The aims of this study were firstly, to determine the presence of ChHV5 in both tumour exhibiting and tumour-free turtles using molecular techniques and secondly, to determine the phylogeography of ChHV5 in Sabah. We also aim to provide evidence of ChHV5 infection through histopathological examinations. A total of 115 Green turtles were sampled from Mabul Island, Sabah. We observed three Green turtles that exhibited FP tumours and were positive for ChHV5.In addition, six clinically healthy turtles were also positive for the virus based on Polymerase Chain Reaction of three viral genes (Capsid protein gene UL18, Glycoprotein H gene UL22 and Glycoprotein B gene UL27). The prevalence of the ChHV5 was 5.22% in asymptomatic Green turtles. Epidermal intranuclear inclusions were identified in tumour lesions upon histopathological examination. Thus, the emergence of ChHV5 in Green turtle in the waters of Sabah could indicate a possible threat to sea turtle populations in the future and requires further monitoring of the populations along the Bornean coast.

## INTRODUCTION

Fibropapillomatosis (FP) is a debilitating neoplastic disease of sea turtles (Alfaro-Núñez et al., 2014; Ene et al., 2015). The first reported case of FP was in Florida in 1938 where a Green turtle was captured with tumour-like characteristics (Smith & Coates 1938). However, it has been reported that disease outbreaks in the wild have increased since the 1980s (Patrício et al., 2016). In Asia, it was reported that FP tumour was first observed in nesting Green turtles (*Chelonia mydas)* from Sarawak Turtle Islands in 1958, but there have been no further reports of possible etiological agents in the waters of Malaysia (Li et al., 2017). However, a high prevalence of FP was observed in Indonesia, a neighbouring country to Malaysia (Adnyana et al., 1997). In addition, FP has been be classified as a pandemic disease which appears to be increasing in some parts of the world (Work et al., 2009). This further serves a need to determine the presence of FP and to better understand how it might affect turtle population in Borneo, a region where the health of sea turtles is poorly understood.

FP is described as a neoplastic conditions where it is clinically characterized by tumour formation in both external (skin) and internally (organs) (Work et al., 2004; Work et al., 2009). The tumour primarily occurs on the eyes, soft tissues, flippers, carapace, plastron, and internal organs and the size may reach 20 cm in diameter (Reséndiz et al., 2016; Li et al., 2017). The affected sea turtles become thin and weak and hamper their daily activities such as feeding and locomotion as the tumour begins to impede organ functions. Eventually the tumours grow large enough and impair buoyancy of the turtle and in several cases causing death (Work et al., 2009; Page-Karjian et al., 2012; Patrício et al., 2016; Reséndiz et al., 2016).

As severe tumour loads could be life threatening to the sea turtles, FP has emerged as an important panzootic disease that needs to be monitored (Page-Karjian et al., 2012; Jones et al., 2016). While the cofactor of the outbreaks from Hawaii, Florida and Indonesia has been hypothesized to be environmental factors, metazoan parasites and oncogenic viruses, the etiologic agent most consistently associated with FP is Chelonid herpesvirus 5 (ChHV5), an alphaherpesvirus from the family Herpesviridae (Klein et al., 1998; Quackenbush et al., 2001; Work et al., 2004; Page-Karjian et al., 2012; Alfaro-Núñez et al., 2016; Li et al., 2017). As such, a thorough understanding of the cause and pathogenesis is needed to aid in controlling efforts against the disease (Page-Karjian et al., 2012).

ChHV5 infections, as with most other Herpes virus infections, result in latent infections that persist overtime. Even though the virus is dormant, the virus transmission is reactivated under certain circumstances such as temperature, environmental change and efficacy of the host’s immune system (Page-Karjian et al., 2012; Jones et al., 2016; Alfaro-Núñez et al., 2016; Li et al., 2017). Consequently, infected sea turtles may be carrying segments of the viral DNA without exhibiting any signs of tumours in the initial stages of infection (Alfaro-Núñez & Gilbert 2014; Patrício et al., 2016).This implies that environmental factors and other putative causes enhance the virus outbreak and facilitates transmission. At times, tumours develop large enough to cause death (Herbst & Klein, 1995; Patrício et al., 2016).

The complete genome of ChHV5 was sequenced using BAC-derived sequences and is known to have a type D genome (Ackermann et al., 2012). Several studies have used classic molecular approaches to detect ChHV5 DNA via polymerase chain reaction (PCR) in tumour samples from turtles affected with FP. Replication of the Herpes-viruses is common in tissues which exhibit tumours as large concentrations of viral particles exist. Hence, the expression of ChHV5 in these turtles is predictable. However, during latent infections, the concentrations of viral particles are very low, presenting a challenge to detect the virus (Alfaro-Núñez et al., 2016; Patrício et al., 2016).

Even though, FP is widely distributed, reports on FP in Asia are few and limited (Hargrove et al. 2015; Li et al., 2017). The identification of ChHV5 in the pristine waters of Sabah’s east coast is important as the site is ecologically rich and home to many marine species. Anecdotal reports indicate that several Green turtles have been spotted with FP tumours. Through this study, we have verified the claims with the aim to document the presence of FP in Green turtles in this region using microscopy and molecular approach. To the best of our knowledge, we present here the first record of ChHV5 in Green turtles from Borneo.

## MATERIAL AND METHODS

### Field sampling

The field sampling was carried out at Mabul Island (4°14′45″N 118°37′52″E) located at the east coast of Sabah, Malaysia, in May and November 2015, and again in November 2016 (Figure 1). Each field trip was conducted for four days, with a total of 13 dives per trip. The study site is a feeding ground for both the Green and Hawksbill (*Eretmochelys imbricata*) turtles. The turtles were caught by hand while SCUBA diving during the day at the seven established dive sites (Figure 1) at depths not exceeding 20 m. The turtles were photographed, measured and tagged on board the research vessel. Two individually numbered Inconel tags from Sabah Parks, Malaysia were applied to the axial scale of each front flipper (Limpus, 1992a, 1992b).

**Figure 1:**
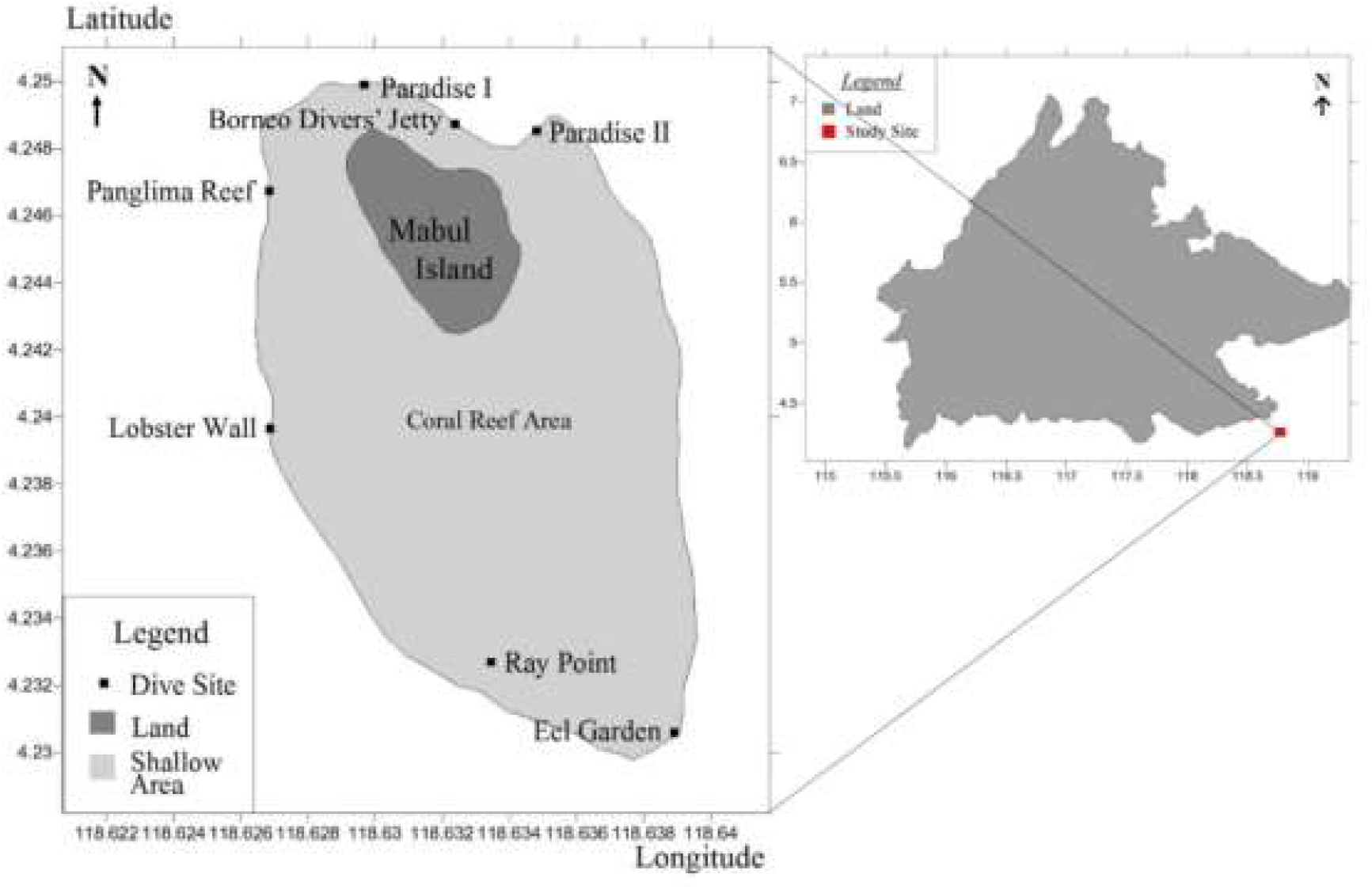
Map of Mabul Island (left), showing the established dive sites where sea turtles were captured, located in south-eastern Sabah (right), Malaysia. (Credit: Haziq Harith Abd Hamid)

The turtles were measured using standard procedures (Limpus & Reed, 1985; Bolten, 1999) and all curved carapace length measurements were taken using a flexible tape to the nearest mm. Tissue samples (n=131) from 115 Green turtles and 16 Hawksbills were obtained from the fore flipper using a 6mm skin biopsy punch after disinfection with Betadine, following the method described by (Dutton, 1996) and stored in lysis buffer. The turtles were also evaluated physically immediately after capture. All 131 samples were subjected for molecular analysis while three of the turtles with tumours were proceeded for pathological analysis. The turtles were then released into the sea after all measurements and tissues were collected. This study was conducted with the permission of the Sabah Wildlife Department.

### Histopathological Examinations

Tissue samples from the tumours (n=3) were fixed in a 10% buffered formaldehyde solution prior to fixation and processed for routine histopathology (Bancroft et al., 2012) and TEM examination (Hunter 1993). The tissues were then embedded in paraffin wax. The paraffin-embedded samples were then sectioned to 4 μm and finally stained by hematoxylin and eosin (HE) using standard techniques for histopathological examination. The slide was viewed using light microscopy to confirm the presence of viral particles in the tumour and characterize the tumour morphologically (Klein et al., 1998; Herbst et al., 1999; Li et al., 2017).

### Molecular analysis

DNA was extracted from 25 mg of tissue using a modified cetyltrimethylammonium bromide (CTAB) method according to the protocol by Bruford et al. (1992). The final purified genomic DNA was eluted in 30 μl of EB buffer. PCR assay was performed with four independent primer sets developed by Alfaro-Núñez et al. (2014) and Lu et al. (2000) for CFHV detection (Table 1). The primers amplified the highly conserved region of the ChHV5 gene (Capsid protein gene UL18, Glycoprotein H gene UL22, Glycoprotein B gene UL27(Alfaro-Núñez et al.,2014) and DNA polymerase catalytic subunit pol UL30 (Lu et al., 2000), the latter have been found to be more sensitive than the other available oligonucleotide primers shown in a previous study (Alfaro-Núñez et al. 2014).

**Table 1:**
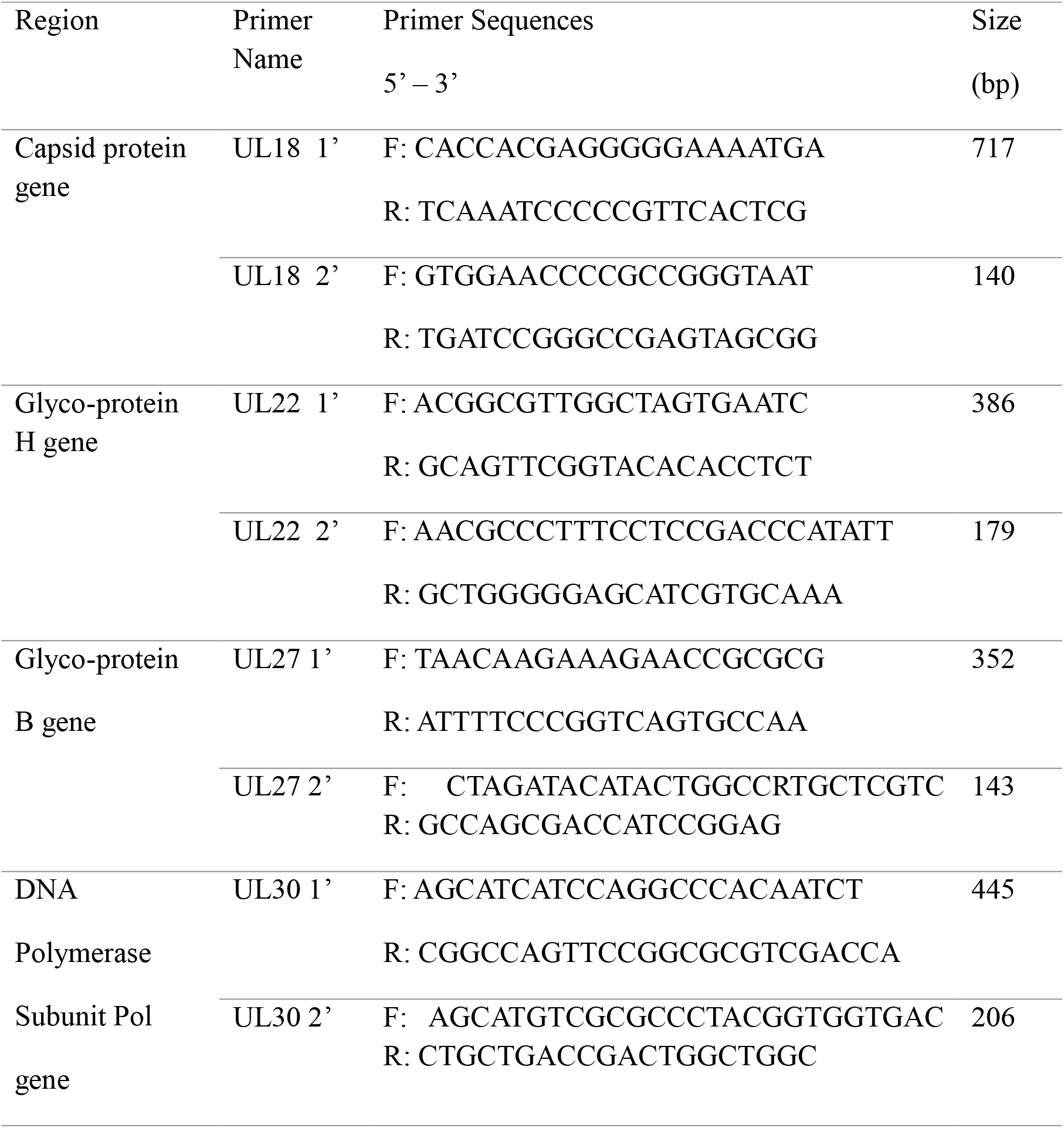
List of nested primers used to amplify the four ChHV5 gene regions and the expected amplification size (bp).

Amplification of the ChHV5 gene regions were carried out set up in 10 µl reaction volume containing 25–30 ng of genomic DNA, 0.2 U GoTaq DNA polymerase (Promega), 1x GoTaq buffer (Promega), 2.0 mM of MgCl_2_, 10 pmol of each primer and 0.4 mM of dNTPs in a Thermocycler (PT1000, Bio-Rad Laboratories). The amplification temperature profiles consisted of an initial denaturation at 94°C for 60 s, followed by 35 cycles of denaturation at 94°C for 30 s, an optimal annealing temperature for 30 s, extension at 72°C for 30 s, and a final elongation step at 72°C for 7 min. The PCR products were visualised on a 1% Agarose gel prior to DNA sequencing.

Positive viral amplicons that were identified were further confirmed by Sanger sequencing using the BigDye Terminator Kit v3.1 and analysed on an ABI 3130 DNA sequencer (Applied Biosystems, Inc.). Viral sequences were compared to the Green turtle Herpes virus sequences available in Genbank to verify gene targets according to DNA identity. This was done through multiple sequence alignment by using the ClustalW program implemented in MEGA5.2. Phylogenetic trees were generated using the BEAST v2.50 software with 10,000,000 heated chain generations and a burn-in fraction of 10% (Bouckaert et al., 2019). FigTree v.1.44 was used to illustrate the phylogenetic tree (Rambaut, 2018).

## RESULTS AND DISCUSSION

We sampled a total of 131 living turtles in the waters of Mabul Island over a period of one and a half years (May 2015 to November 2016). The island, along with its well-known neighbour Sipadan Island, is a foraging ground for Green and Hawksbill turtles. Most of the turtles were deemed clinically healthy with overall good body condition (convex-shaped plastrons) and size. Green turtles were mainly juveniles with curved carapace lengths ranging from 524 mm to 840 mm and more abundant than Hawksbills (1 Hawksbill: 7 Green turtle ratio.

Three Green turtles with visible FP tumours were encountered at Mabul Island. The tumour scores were defined based on the study proposed by Work & Balazs (1999) where the tumours were classified into four categories. Size A (< 1 cm) Size B (1 – 4 cm), Size C (4 – 10 cm) and finally Size D (> 10 cm). In this study, the tumours were comprised of all four categories. The largest tumour observed was 12 cm in size which was present on the ventral surface of the turtle. Greenbaltt et al. (2005), suggested that the variation in size and morphology of the tumours may depend on geographical regions and where some parts of the world may have a higher percentage of corneal FP while other parts may only have epidermal tumours.

We observed that morphologically all the tumours were a combination of either smooth or verrucous. Most tumours were soft while some appeared to be rigid. The nodules were firm and pale, with similar morphology to those described by Greenbaltt et al. (2005) and Work et al. (2004). One individual had a debilitating advanced stage of FP with tumours in all regions of the body (hind flippers, plastron, neck and carapace) (Figures 2a & 2b). Multiple nodules, ranging from 1 cm to 7 cm in diameter, were present on the ventral surfaces of both back flippers. The nodules were filled with parasitic leeches. However, these leeches were not further examined. The other two turtles had tumours on the eye lid (Figure 2c) which impacted their vision.

**Figure 2a:**
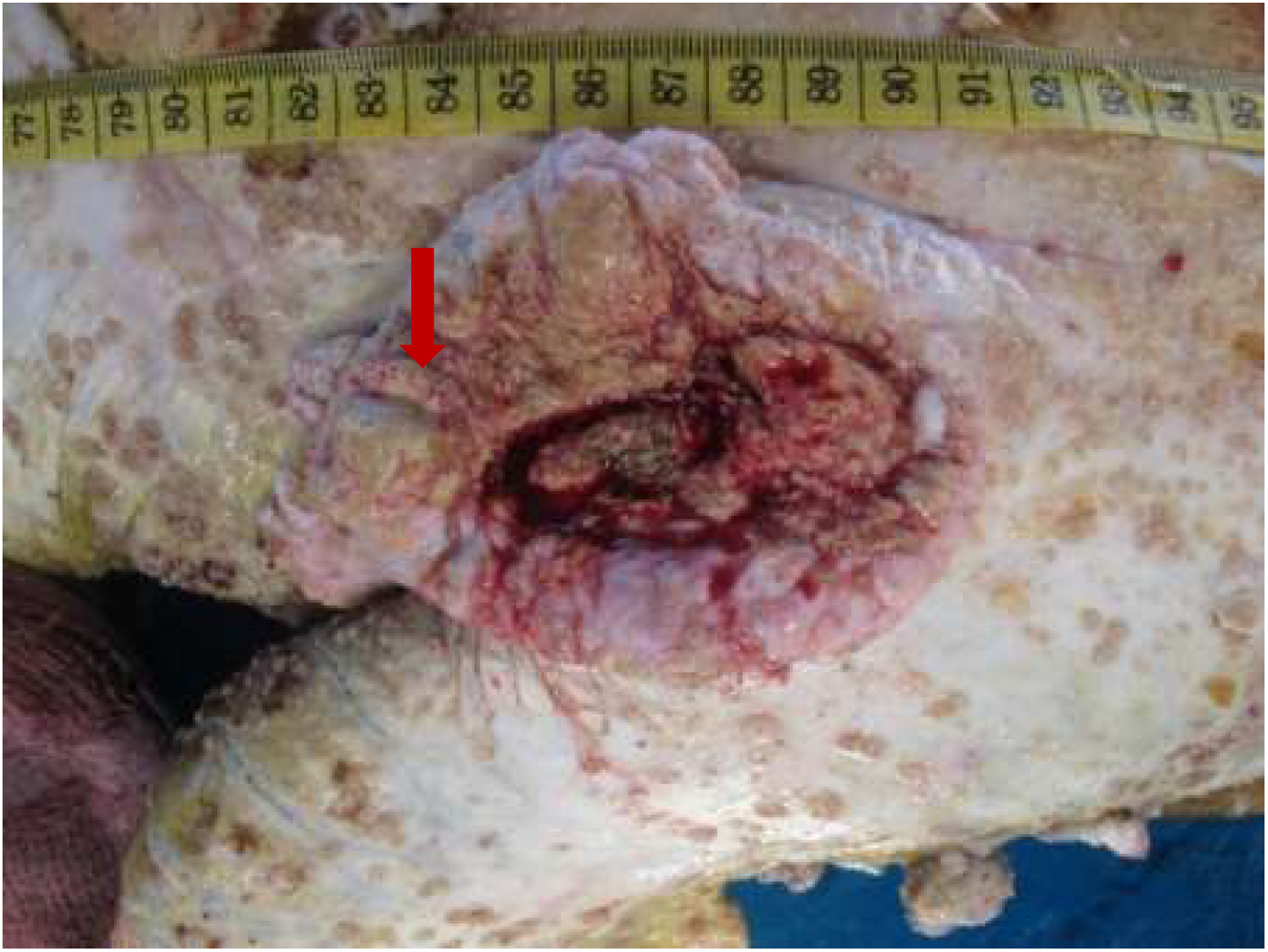
A 12 cm tumour (arrow) present on the ventral surface of a Green Turtle.

**Figure 2b:**
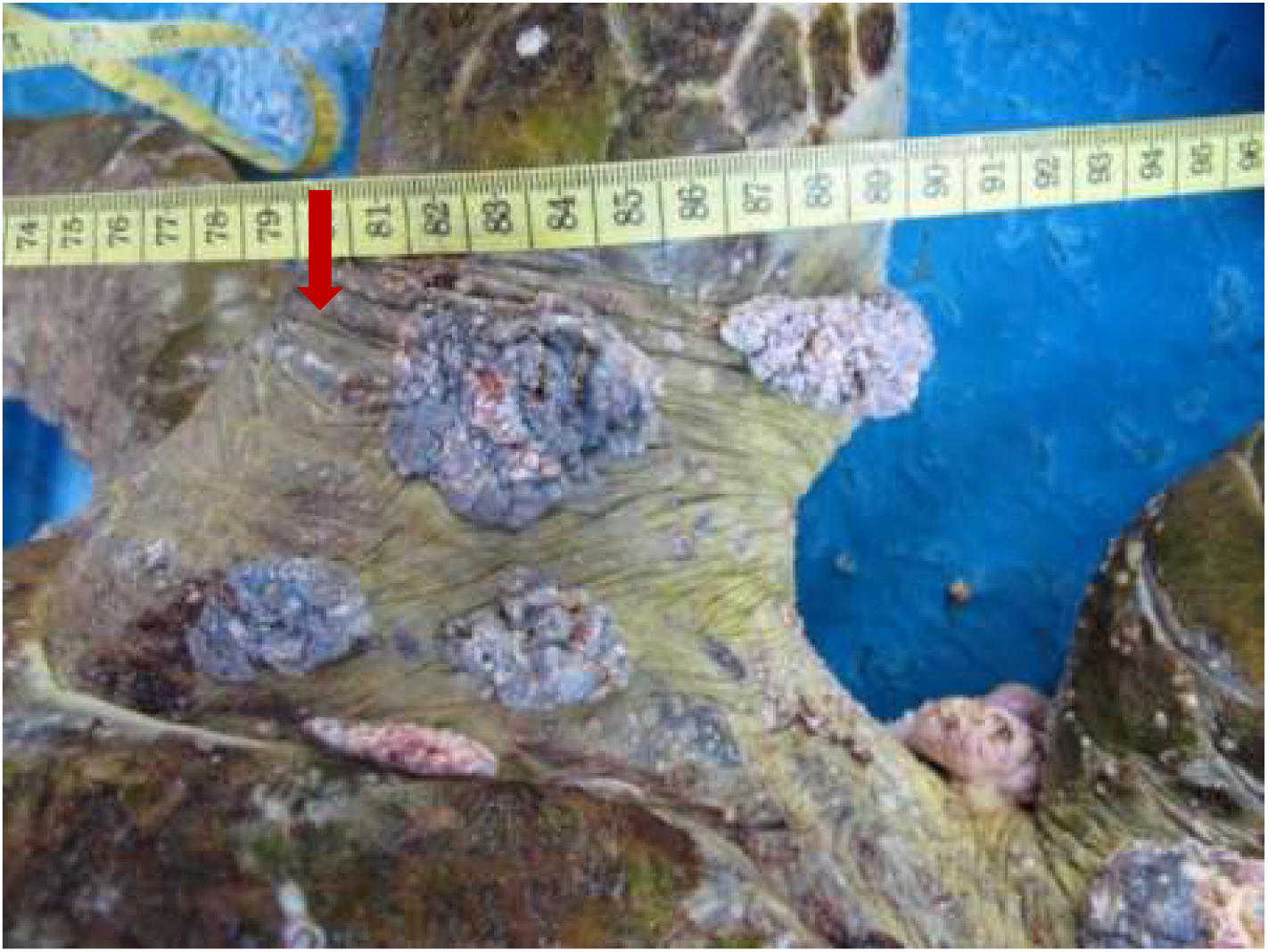
Multiple verrucous nodules (arrow) on the dorsal surface of the hind flipper of a Green turtle.

**Figure 2c:**
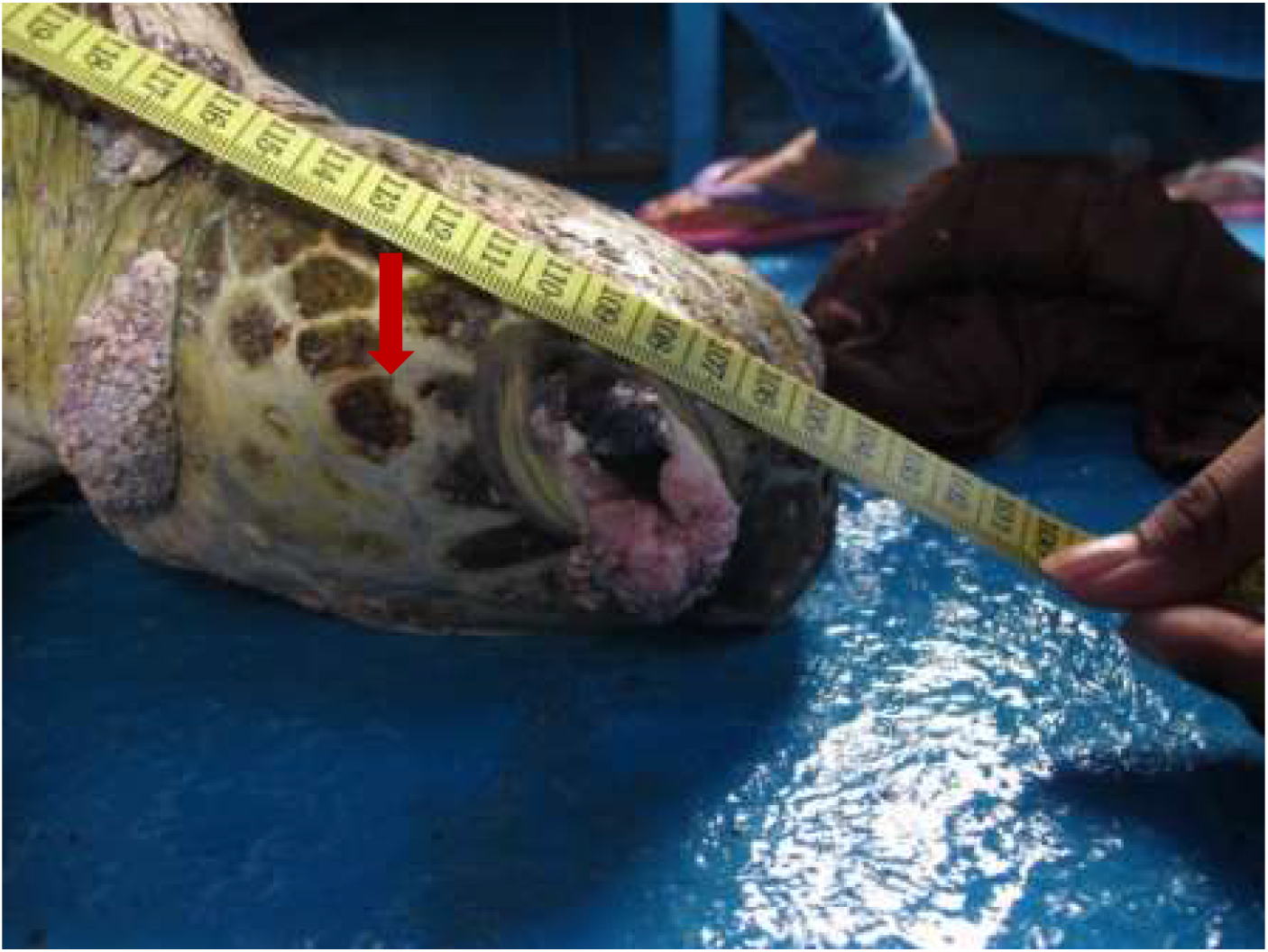
A 4 cm verrucous lesion at the lateral canthus of the eye of a Green turtle.

Histopathological examinations were conducted to provide evidence of ChHV5 in tumour tissue. Light microscopic examination of the tumour lesion that was biopsied revealed the presence of Horn Cyst (Figure 3). Hyperkeratotic epidermis (characterized as papilloma) and acanthosis with Epidermal intracellular inclusion bodies (EII’s) was also identified (Figure 3a–c). The presence of Horn Cyst, which classically indicatives the presence of a viral infection in tumours further confirms ChHV5 infection in the tumoured tissues (Wang et al., 2007). The Hyperkeratotic epidermis and acanthosis with Epidermal intracellular inclusion bodies (EII’s) implies a transcriptionally active (i.e., non-latent) state of ChHV5 in FP tumours of the affected Green turtles. Work et al. (2014) stated that IILs are commonly found on the outer layer of the epidermis. Therefore, it is transmitted easily when in contact with other foraging turtles or spread into the environment. This observation is consistent with the previous reports for FP in sea turtles by Brenes-Chaves et al.(2013), Monezi et al.(2016) and Work et al.(2014), and further supports the initial hypothesis by Jacobson et al., (1989) that the virus is replicating in FP tumours from sea turtles. The results seem to indicate that ChHV5 plays an active role in FP of Green turtles and could be horizontally transmitted between regions based on the phylogenetic analysis.

**Figure 3a:**
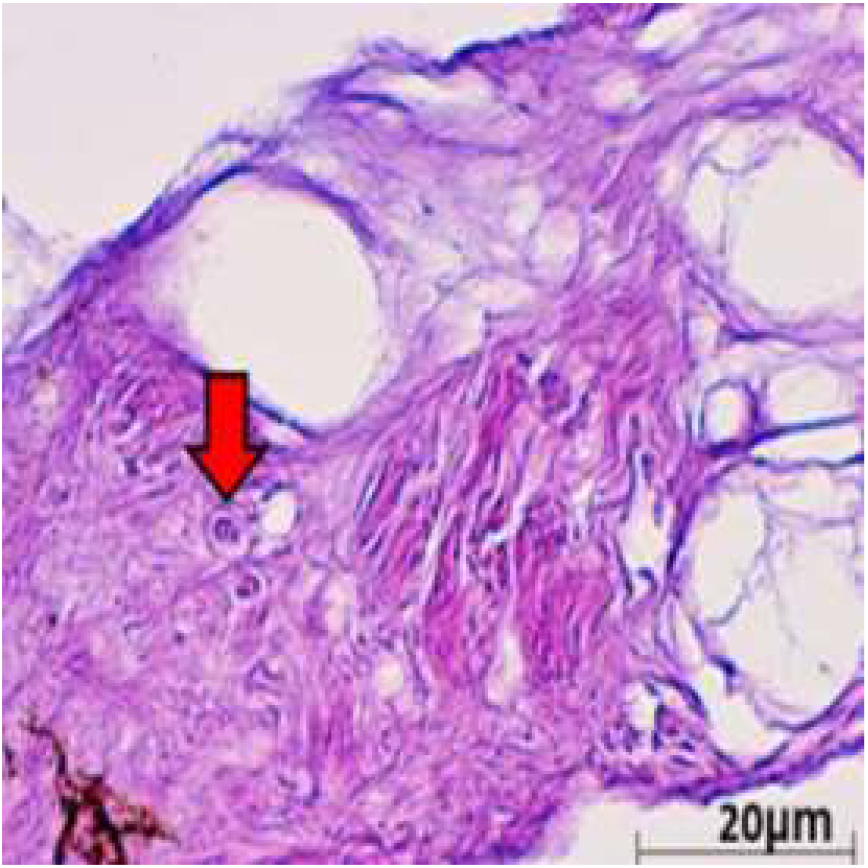
Histological cross section of one of the verrucous nodulations sampled from the ventral surface of the hind flipper of a Green turtle where it shows epidermal internuclear inclusions (arrow) within epidermis manifesting ballooning degenerations. (40× magnification with hematoxylin and eosin stain; scale bar = 20 μm).

**Figure 3b:**
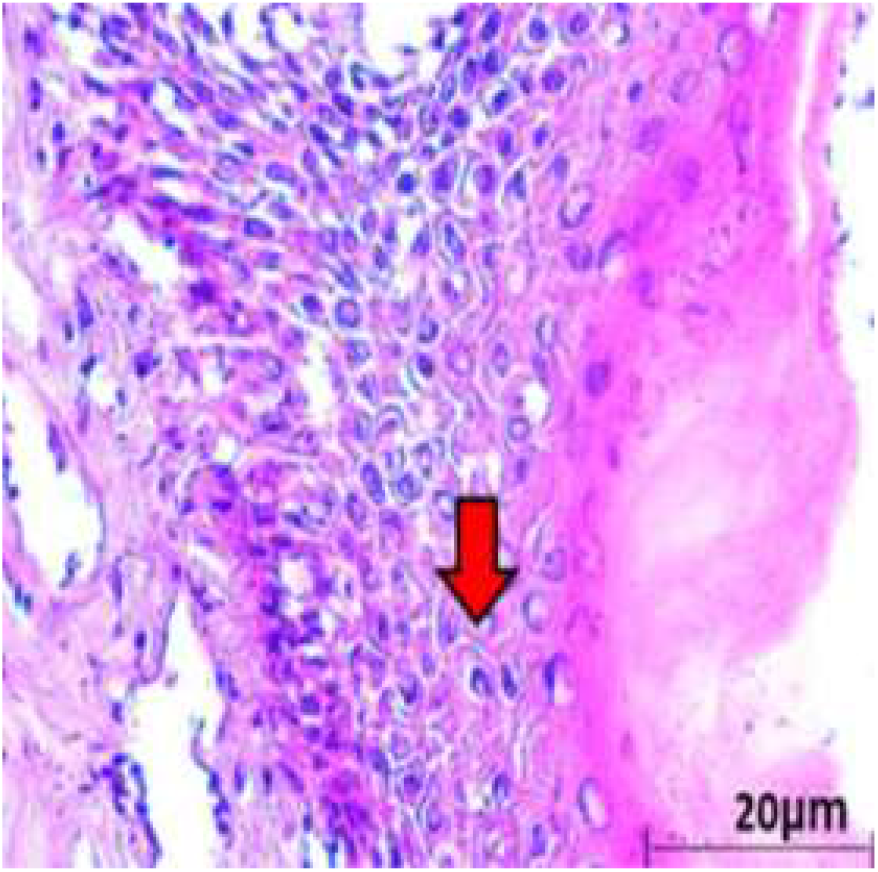
Histological cross section of one of the verrucous nodulations sampled from a Green turtle where it shows Acanthosis and orthokeratotic hyperkeratosis with necrosis of the stratum basale (arrow). (40× magnification with hematoxylin and eosin stain; scale bar = 20 μm).

**Figure 3c:**
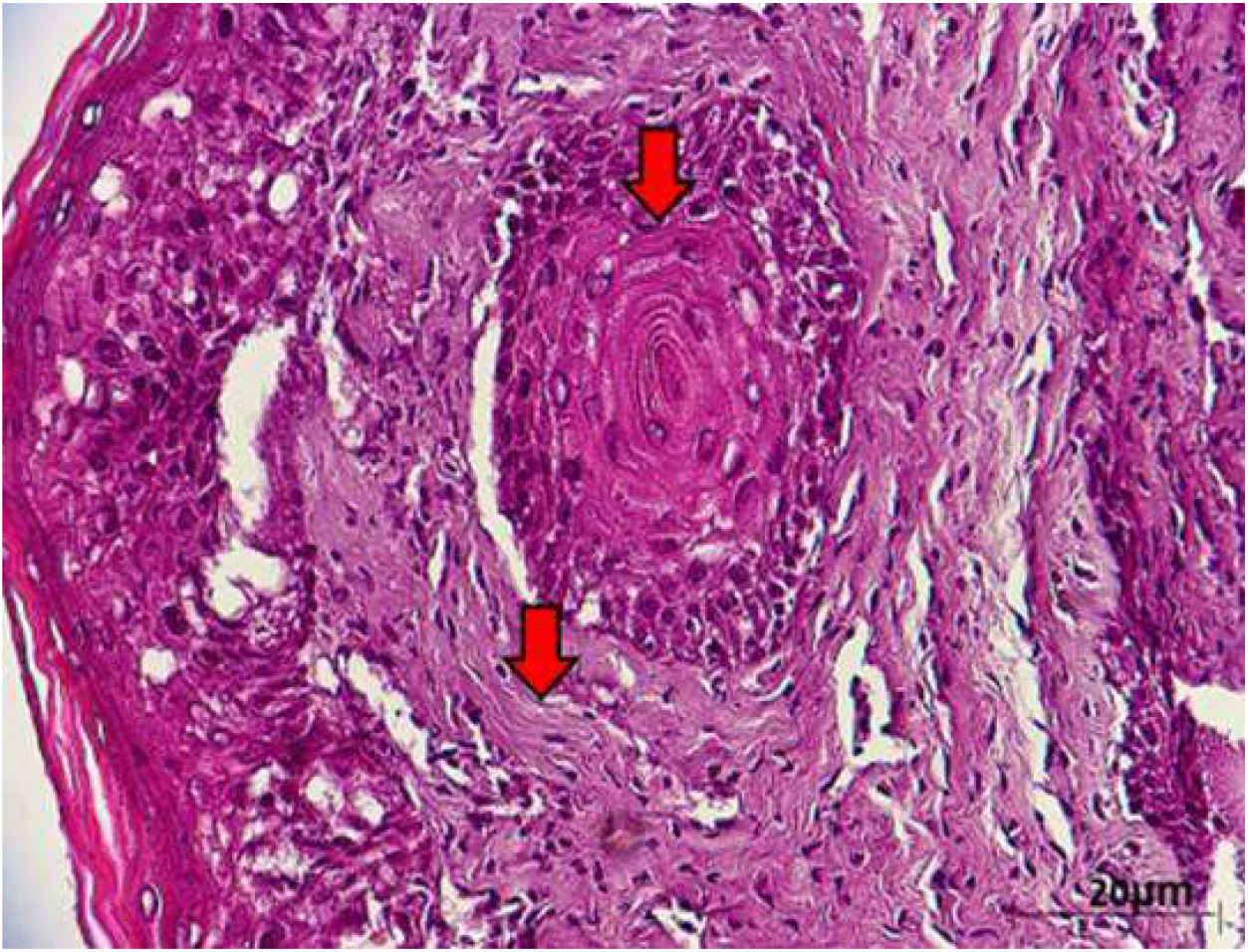
Histological cross section of one of the verrucous nodulations sampled from the ventral surface of the hind flipper of a Green turtle where it shows epidermal hyperplasia (arrow) (40× magnification with hematoxylin and eosin stain; scale bar = 20 μm).

**Figure 3d:**
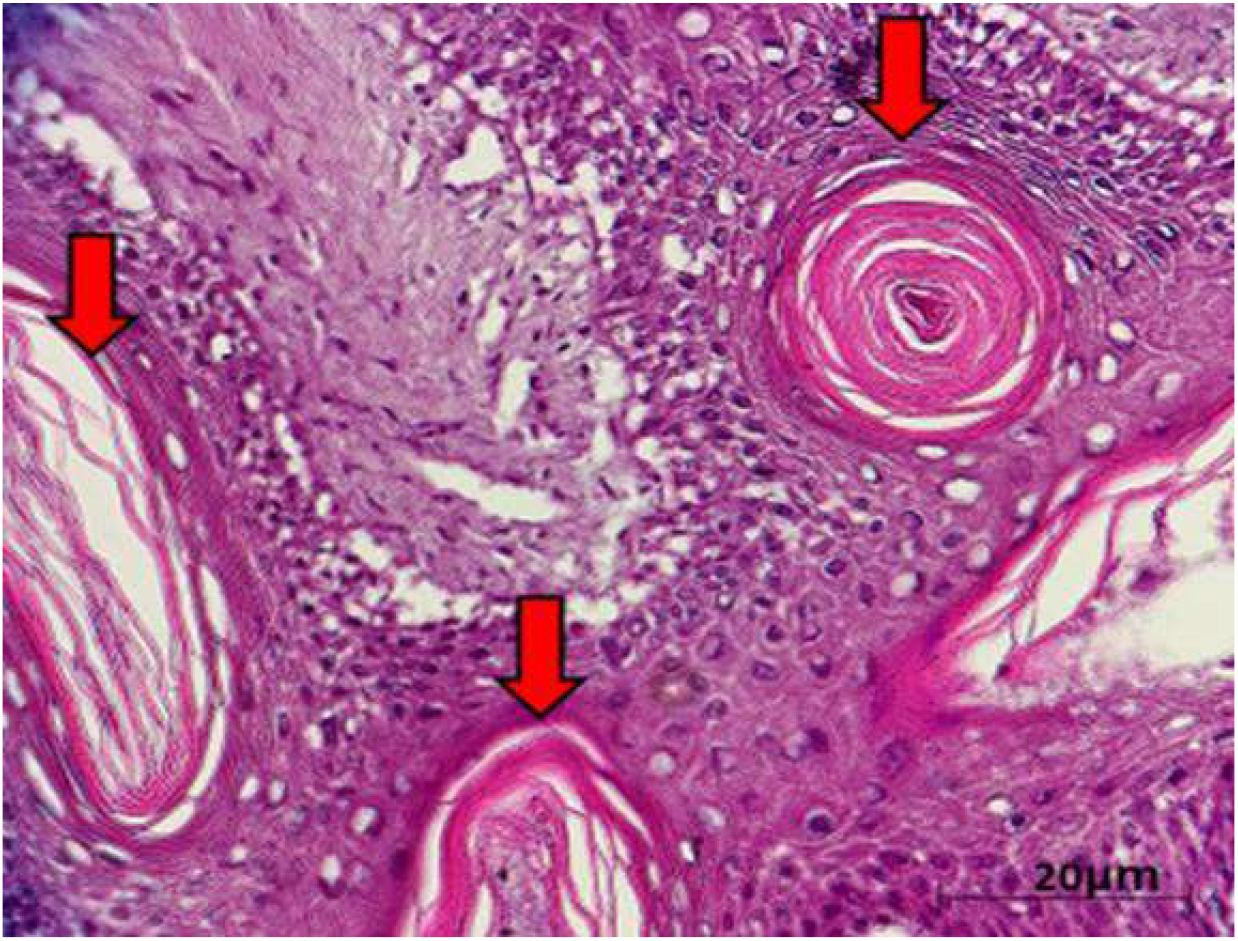
Histological cross section of one of the verrucous nodulations sampled from the ventral surface of a Green turtle where it shows horn cyst (arrow) (40× magnification with hematoxylin and eosin stain; scale bar = 20 μm).

In addition to the gross and histopathological examinations, molecular analysis was conducted to provide additional evidence of ChHV5 infection. A total of 131 (115 Green and 16 Hawksbill) individual sea turtles were screened for ChHV5 (including the three with visible signs of tumour) using PCR by amplifying sequences that represent the four viral genes (Capsid protein gene UL18, Glycoprotein H gene UL22, Glycoprotein B gene UL27 and DNA polymerase catalytic subunit pol UL30). All 16 Hawksbill turtle tested negative for the ChHV5 virus. Of the 115 Green turtles analysed (FP-exhibiting and clinically healthy), nine Green turtles (3 FP exhibiting and 6 clinically healthy turtles) were PCR-positive for ChHV5 (Table 2). Four samples were positive for the Capsid protein gene UL18, nine samples were positive for the Glycoprotein H UL22 gene while six had sequences of the Glycoprotein B gene UL27. As for the DNA Polymerase gene UL30, five positive samples were detected. A previous study by Alfaro-Nunez & Gilbert (2014) concluded that, even though the sea turtles are infected with ChHV5 virus, it is very unlikely to indicate PCR-positive if only a single gene is used to screen for the presence of the virus. Consequently, the combination of all the four genes allowed the screening of ChHV5 positive samples with differing stages of infection.

**Table 2a:**
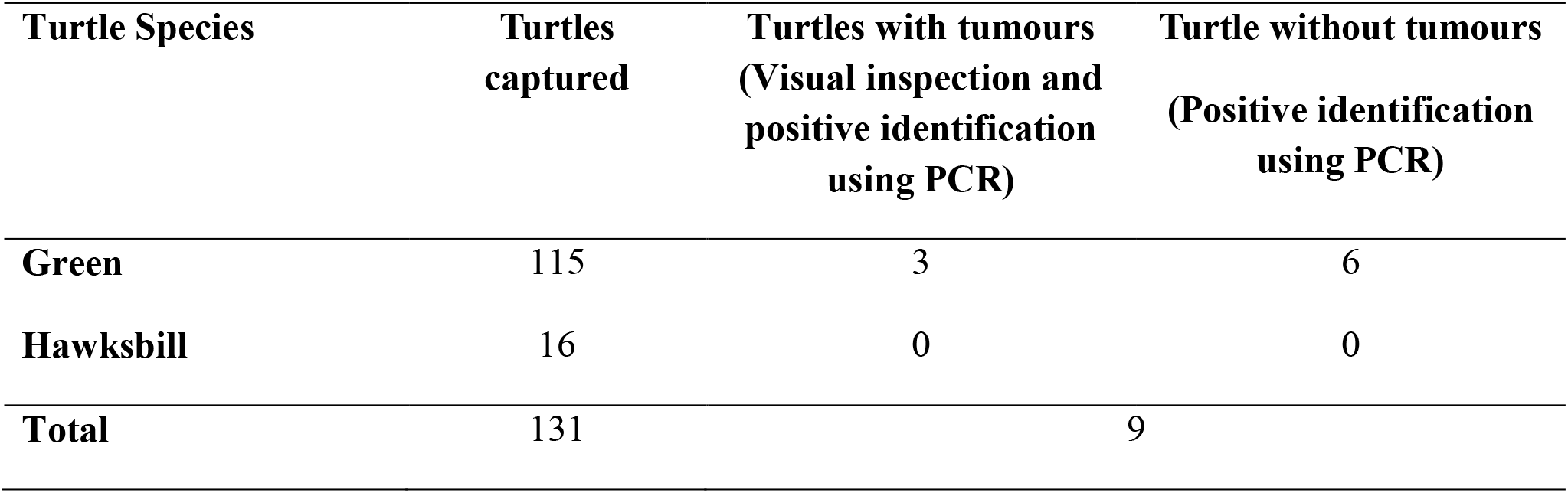
Detection of ChHV5 in tissues with and without tumours in the Green and Hawksbill turtles.

**Table 2b:** Sample was identified positive if any of the four viral genes showed amplification and positively verified through DNA sequencing. There was no presence of ChHV5 detected in the Hawksbill turtles.

| Gene | Green Turtle ID |  |  |  |  |  |  |  |  |
| --- | --- | --- | --- | --- | --- | --- | --- | --- | --- |
|  | FP8* | FP41 | FP45 | FP46 | FP48 | FP50 | FP70* | FP71* | FP115 |
| **Capsid Protein (UL18)** | √ |  |  | √ | √ |  | √ |  |  |
| **Glycoprotein H (UL22)** | √ | √ | √ | √ | √ | √ | √ | √ | √ |
| **Glycoprotein B (UL27)** | √ | √ | √ | √ |  | √ |  | √ |  |
| **DNA Polymerase (UL30)** | √ | √ | √ |  | √ |  |  |  | √ |

Note: \* Turtle with tumours (Visual inspection and positive identification using PCR)

Various theories have been suggested for ChHV5 to be present in both FP-exhibiting and clinically healthy turtles. At present, latent infections is the most accepted theory for these conditions. Like most Herpesvirus infection characteristics, the virus is capable to cause latent infections in their host (Lu et al., 2000; Greenblatt et al., 2004). This characteristic plays a crucial role during latent infections, as the viral gene is still present but the expression is minimised. Albeit using four genes for the detection of ChHV5, Alfaro-Núñez & Gilbert (2014) hypothesized that the detection accuracy is proportional to the viral DNA concentration and the DNA fragment sizes of the tumour sample. Similarly, the ability of the viral DNA to anneal to the with the PCR chemistry plays a major factor as well. As such, it present a challenge in detecting ChHV5 in latent infections. Hence, an absence of tumour does not necessarily indicate the absence of ChHV5 infection. This theory was supported by previous research conducted by Alfaro-Nunez et al. (2014) and Jones et al. (2016).

All ChHV5 positive sequences from this study have been deposited in Genbank (https://www.ncbi.nlm.nih.gov/) with the accession numbers: MG894341-MG894365. Phylogenetic analysis inferred using the ML method showed clusters and distribution of the ChHV5 strain from Mabul Island and isolates from other geographical regions. The phylogenetic tree was constructed from the sequences obtained from BLAST nucleotide search. The phylogenetic tree were constructed with capsid protein gene (UL18), glycoprotein H gene (UL22), glycoprotein B (UL27) and DNA polymerase gene (UL30) sequences (Figure 4a–d).

**Figure 4a:**
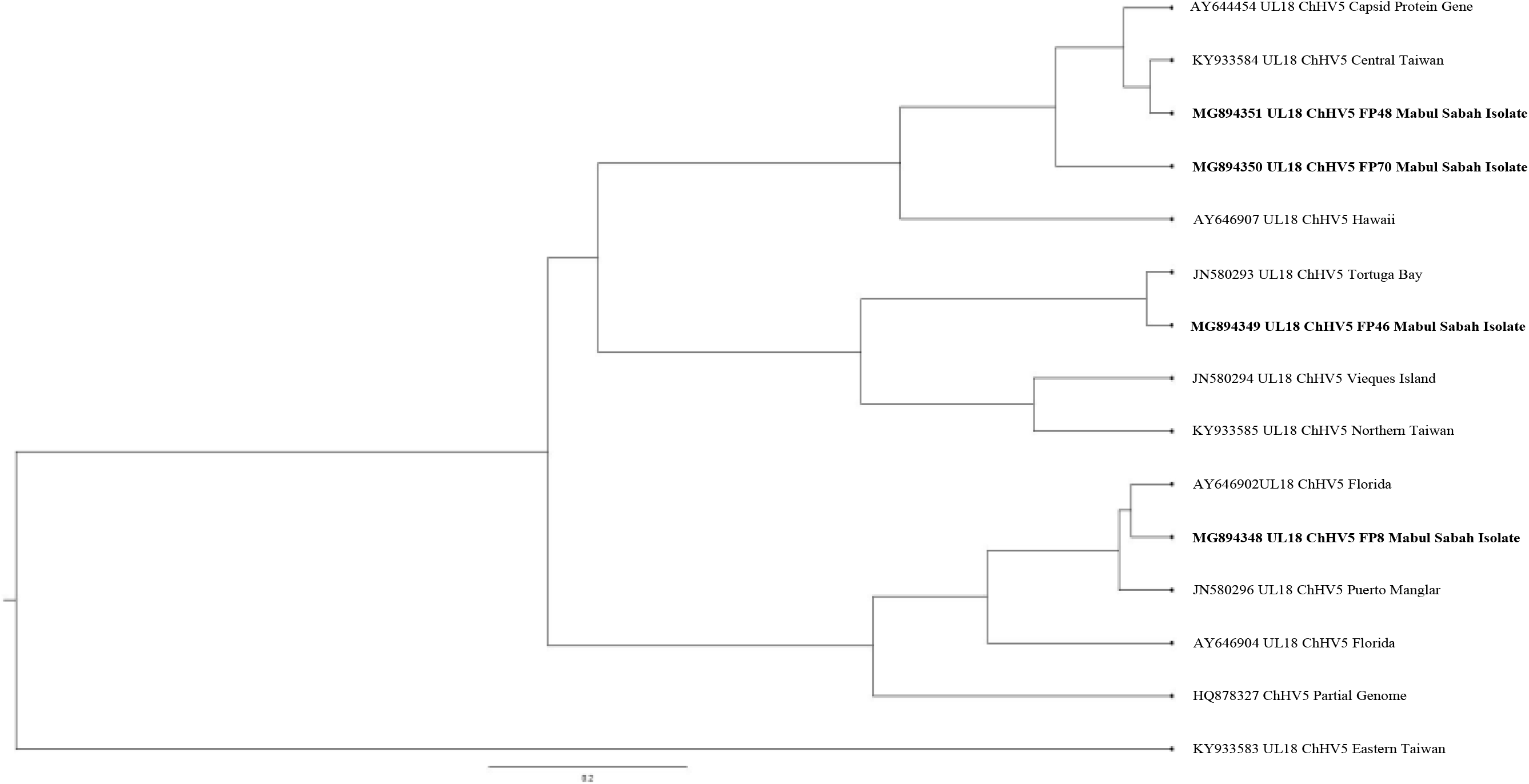
Cluster analysis and distribution of ChHV5 variants inferred using BEAST v2.50. The ChHV5 Major capsid protein gene (UL18) from Mabul Island indicates that the Mabul Island strain is clustered closely to the Brazialian strain of Vieques Island, Puerto Manglar and Tortuga Bay.

**Figure 4b:**
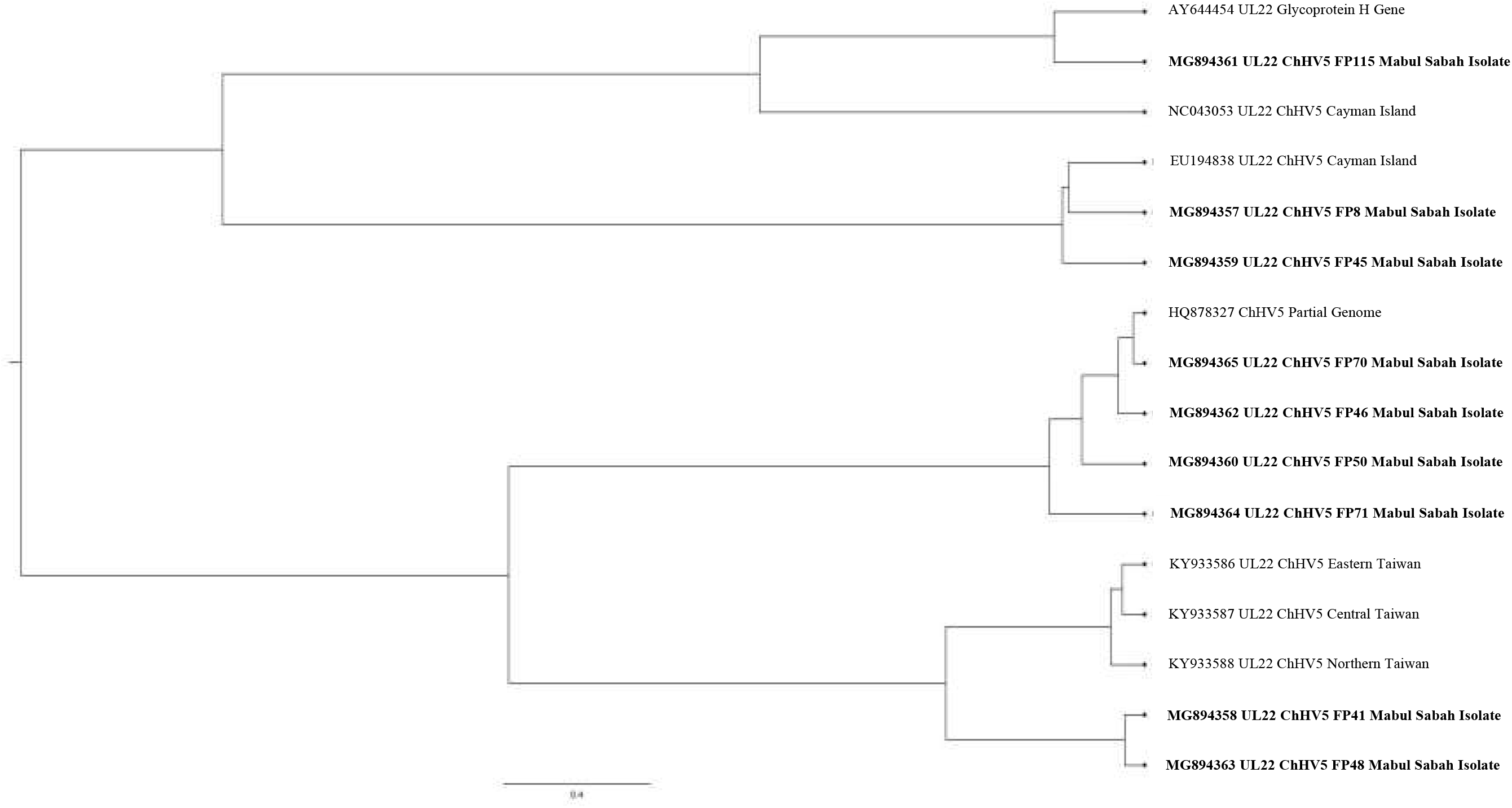
Cluster analysis and distribution of ChHV5 variants inferred using BEAST v2.50. The ChHV5 glycoprotein H gene sequences (UL22) indicate that the Mabul Island strain is clustered together with those from Cayman Island. Four viral sequence of the ChHV5 glycoprotein H gene sequences (UL22) from Mabul Island strain were clustered into a separate branch.

**Figure 4c:**
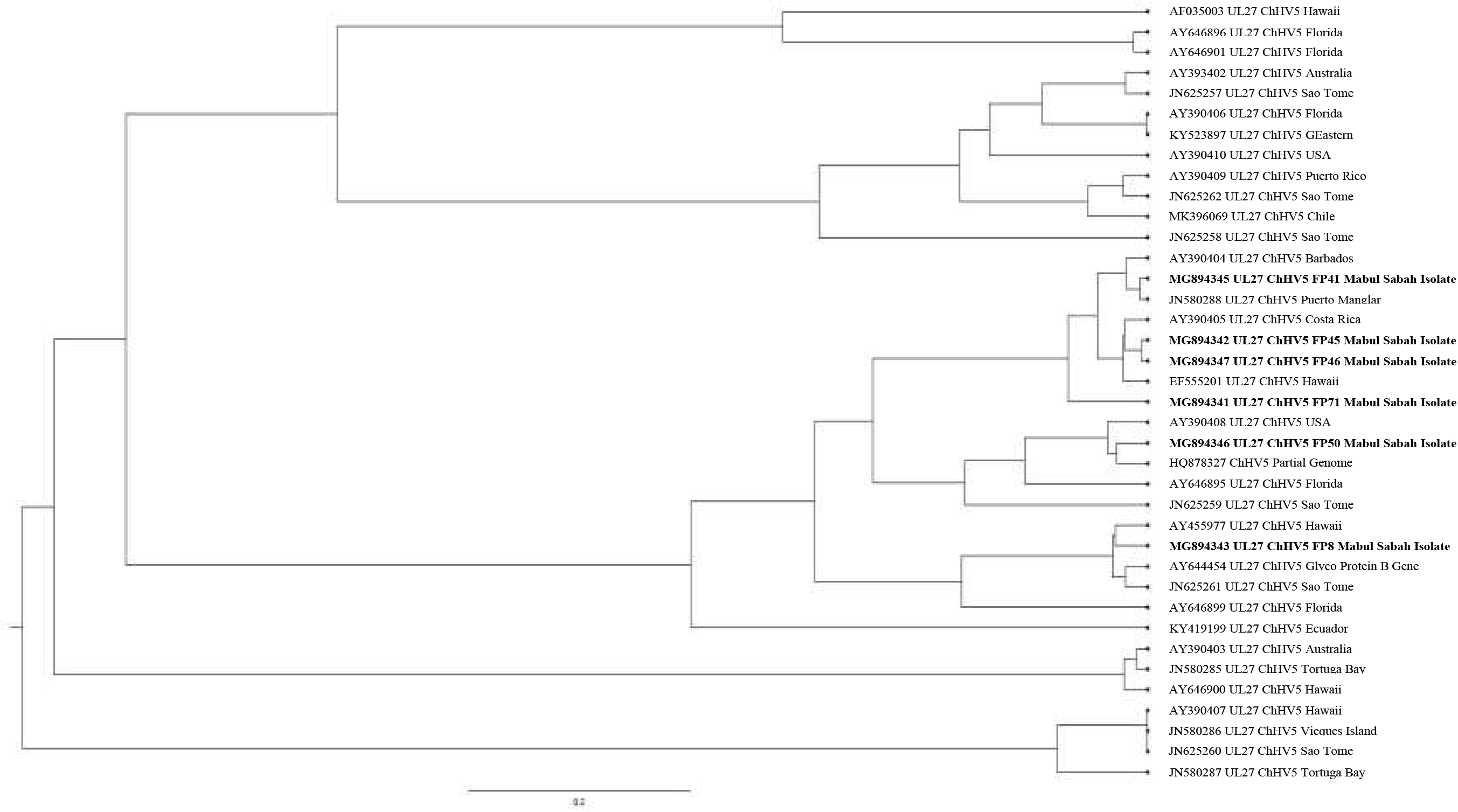
Cluster analysis and distribution of ChHV5 variants inferred using BEAST v2.50 The Glycoprotein B (UL27) gene sequences indicates that the Mabul Island strain have more similarity to the Hawaiian and Brazilian strain.

**Figure 4d:**
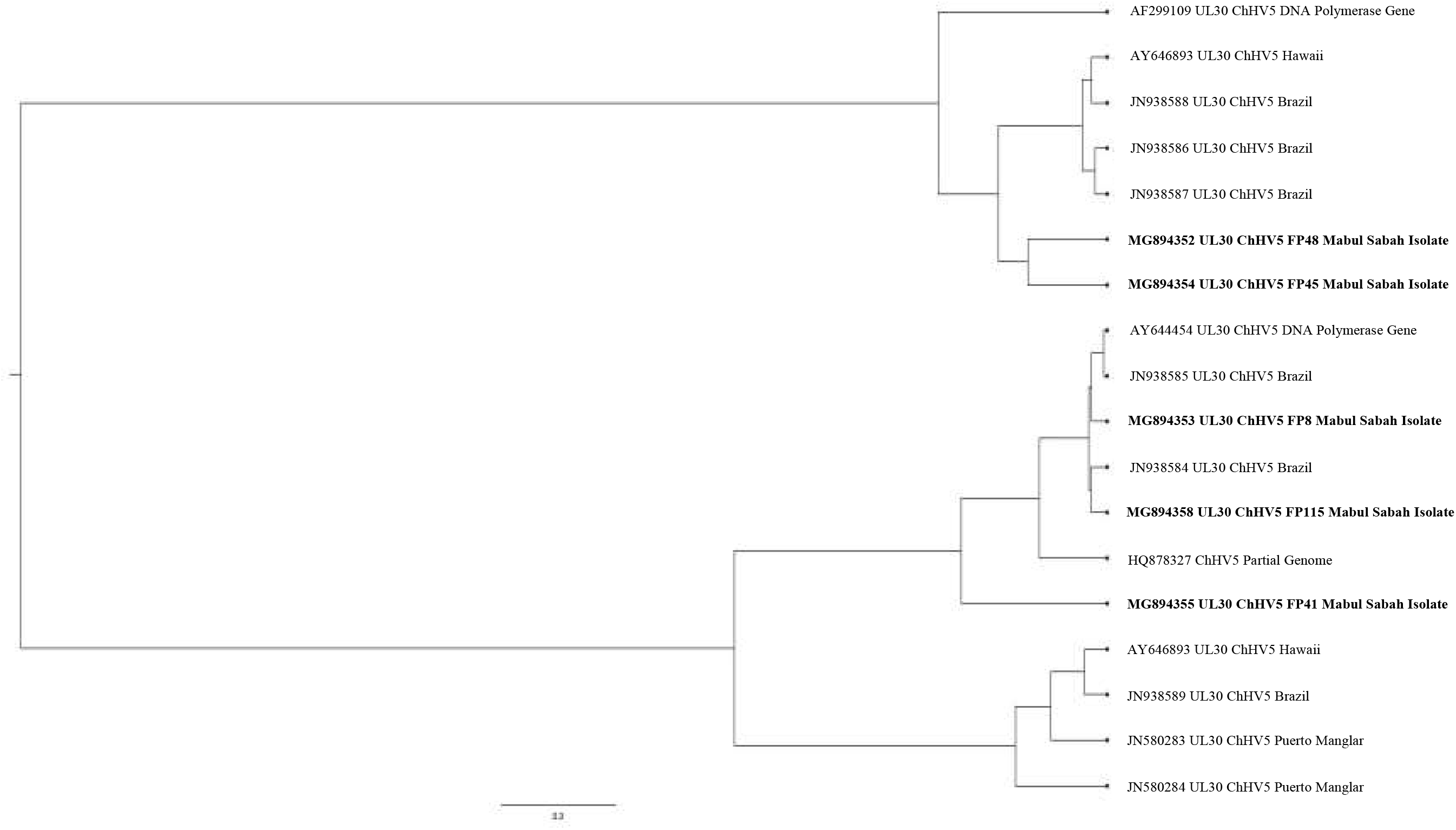
Cluster analysis and distribution of ChHV5 variants inferred using BEAST v2.50 Phylogenetic tree of UL30 DNA polymerase gene shows that the ChHV5 variants from Mabul Island are closely related to the variants from Florida.

Phylogenetic analysis inferred using BEAST v2,50 and visualized using FigTree software (http://tree.bio.ed.ac.uk/software/figtree/) showed clusters and distribution of the ChHV5 strain from Mabul Island and isolates from other geographical regions. The ChHV5 Major capsid protein gene (UL18) from Mabul Island indicates that the Mabul Island strain is clustered closely to the Brazialian strain of Vieques Island, Puerto Manglar and Tortuga Bay.

On the other hand, the ChHV5 glycoprotein H gene sequences (UL22) indicate that the Mabul Island strain is clustered together with those from Cayman Island. Four viral sequence of the ChHV5 glycoprotein H gene sequences (UL22) from Mabul Island strain were clustered into a separate branch. Some of the viral sequence of ChHV5 major capsid protein gene (UL18) and ChHV5 glycoprotein h gene (UL22) from Mabul island followed similar pattern, clustering with the viral strain from Taiwan. A recent study by Li et al. (2017) compared Glycoprotein B gene from Taiwan indicated that the Taiwanese strain was also grouped with those from Hawaii, Sao Tome and Puerto Rico which is similar to the Mabul strain.

The Glycoprotein B (UL27) gene sequences have more similarity to the Hawaiian and Brazilian strain. This could be due to interoceanic connection and ad-mixture of population. Although further analysis is needed to support this theory. As for the DNA polymerase gene (UL30), the phylogenetic analysis showed that most of the Mabul Island strain is clustered together with the Brazilian strain.

The general result inferred from the phylogenetic tree may not represent the full genetic distribution based on geographical location as the study is based on a small sample size. However, the phylogenetic tree generally infers the movement of the sea turtle host reflecting if the ChHV5 have undergone region-specific co-evolution through the sea turtle host. The phylogenetic analysis is also important to identify the differences in gross manifestation across different geographical region. For example, there are higher prevalence of oral tumour in Hawaii compared to Florida which have zero cases of oral tumour (Morrison et al., 2018).

As for the Mabul strains, although the ChHV5 viral strain were clustered together with the Hawaii strain, there were no oral tumours found morphologically. Nevertheless, the ability for the virus to cause latent infection, where there is minimum expression of the viral gene should be considered an important aspect. Furthermore, the tumour formation could be induced in specific tissue and absent in other tissues (Alfaro-Nunez et al., 2014). As the phylogenetic result showed that there is no clear distinction based on geographical location, the interaction between host, pathogen and environment should be considered.

A previous study by Greenblatt et al. (2005), implied that sea turtles may have a passive infection where the pathogenesis of the infection depends on environmental triggers such as pollutants and marine leeches. Similarly, Tristan et al. (2010), reported that sea turtle living in habitats that are nearby urban development and are of close proximity with human contact have increasing chances to develop the tumours.

A possible explanation from a previous study is that the sea turtles may have come in contact with environmental pathogens such as parasitic leeches during the migration period (pelagic phase) of the sea turtles, where the turtles are capable of migrating hundreds to thousands of kilometres to different geographical region in search of foraging grounds. These parasitic leeches have been suggested to be the mechanical vector for ChHV5 in Green turtles (Tseng and Cheng, 2013). It was also documented that the viral particle could survive in the ocean for a short period of time before the virus capsid protein degrades (Alfaro-Nunez et al., 2014).

Although, we observed parasitic leeches in the tumours of the turtles, unfortunately we did not perform further examination, which may serve as a limitation of this study. Other co-factors such as environmental (in addition to the parasitic leeches) could contribute and influence the expression of virus. We suggest that future study should include a larger sample size together with the analysis of the parasitic leech to further support this theory.

Herpesviruses are horizontally transmitted from one infected host to another by bodily fluid such as saliva, mucus or direct physical contact (Work et al., 2014; Alfaro-Nunez and Gilbert, 2014). Hence, the virus from the infected turtle may be transmitted to healthy sea turtles. This is made possible as the juvenile and sub-adult sea turtles undergo long distance migrations known as pelagic life stage, migrating between the juvenile nursery habitat and developmental areas while residing there for years (Seminoff and Walace, 2012). Thus, there is a high probability that the sea turtles acquired the infection during this period of their life cycle. Hence, in a foraging ground, the population of sea turtles may be from different geographical regions. This is why only a certain number of the sea turtles in a population develop ChHV5 (Tristan, 2010). This may also explain why the Mabul island ChHV5 viral strain are generally clustered together with different population from different region such as the Taiwan, Brazilian and Hawaian viral strain respectively.

Jones et al. (2016) suggested that the variability in host immunity and geographic regions could influence the viral variants of an infection. To date there are six variants documented for Chelonid herpesvirus (ChHV5) ChHV5 1, 5 and 6 are designated for marine turtles and ChHV5 2, 3 and 4 are described for freshwater turtles. The variants of a virus could have diverse virulence levels and as such the severity and disease presentation may differ with each variant and could become geographically specific.

Furthermore, a study by Patricio et al. (2012), suggested that the movement of the virus (sea turtle host) to these regions could be clarified by the strong equatorial ocean current that supports the flow of the virus. In support of this, Putman and Naro-Maciel (2013), stated that the particles in the ocean follows the movement of TransAtlantic dispersion of sea turtles that involves persistent connectivity between the southwestern Indian Ocean and the South Atlantic. As such, we suggest that a comprehensive study on the migratory routes of the sea turtle population and oceanic current is required to further understand the missing link between geographical connection and ChHV5.

## CONCLUSION

The results of this study facilitate the understanding of the origin and extent of FP in Mabul Island as the disease is considered to be one of the major threats to *Chelonia mydas.* Turtles infected with FP often do not survive as the tumour obstructs vision and impedes daily activities such as feeding and locomotion. This hampers organ functions and eventually causes death. The 5.22% prevalence in asymptomatic green turtles in this study implies that despite living in different geographical regions, Green turtles may be exposed to a common pathogen or other environmental factors that aggravates FP. Even though ChHV5 has evolved for millions of years with its turtle host, for the conservation of these endangered sea turtles, ChHV5 needs to be considered as a re-emerging virus, which threatens Green turtles in marine waters surrounding Borneo and worldwide. In particular, the need to gather more information on sea turtle disease risk analysis and disease hazards from various regions is vital. Identification of the cause of FP will be the first step towards developing effective measures for management and control programs.

## ACKNOWLEDGEMENTS

This study was conducted with the permission of the UMS Animal Ethics Committee and Sabah Wildlife Department. A.L.L acknowledges the MyBrain scholarship from the Ministry of Higher Education, Malaysia. This work was partially supported through the Nagao Natural Environment Foundation (NEF) Grant. The field work in Mabul Island was financially supported by Borneo Divers and Sea Sports (Sabah) Sdn Bhd and the Mabul Turtle Conservation Fund.

